# Collagen density defines 3D migration of CTLs and their consequent cytotoxicity against tumor cells

**DOI:** 10.1101/2021.03.16.435689

**Authors:** Renping Zhao, Xiangda Zhou, Essak S. Khan, Dalia Alansary, Kim S Friedmann, Wenjuan Yang, Eva C. Schwarz, Aránzazu del Campo, Markus Hoth, Bin Qu

## Abstract

Solid tumors are often characterized by condensed extracellular matrix (ECM). The impact of dense ECM on cytotoxic T lymphocytes (CTL) function is not fully understood. Here, we report that CTL-mediated cytotoxicity is substantially impaired in dense collagen matrices. Although the intrinsic killing machinery including expression of cytotoxic proteins and degranulation was intact, CTL motility was substantially compromised in dense collagen. We found that for 3D CTL migration, persistence and velocity was regulated by collagen stiffness and the porosity, respectively. Interestingly, 3D CTL velocity is strongly correlated with their nuclear deformability/flexibility during migration, which is regulated by the microtubule network. Moreover, CTL migration was completely abolished by inhibition of actin polymerization and or myosin IIA. Remarkably, disruption of the microtubule-networks significantly improves the impaired migration, search efficiency, and cytotoxicity of CTLs in dense collagen. Our work suggests the microtubule network as a promising target to rescue impaired CTL killing capacity in solid tumor related scenarios.

## Introduction

Cytotoxic T lymphocytes (CTLs), which are the activated CD8^+^ T cells, compose an essential arm of the immune system to fight aberrant cells like tumorigenic and pathogen-infected cells (Halle et al., 2017; Zhang and Bevan, 2011). CTLs recognize their targets via engagement of T cell receptors (TCRs) with the cognate antigens presented on the surface of target cells (Artyomov et al., 2010; Huang et al., 2007; Li et al., 2013). Once the matching antigens are identified, activation of TCRs triggers the downstream signaling cascades to re-orientate the CTL killing machinery towards the contact site, termed immunological synapse (IS) (Bromley et al., 2001; Dieckmann et al., 2016; Dustin et al., 2010). The major killing mechanisms employed by CTLs are lytic granules (LGs) and Fas/FasL pathway (Barry and Bleackley, 2002). LGs contain cytotoxic proteins such as pore-forming protein perforin and serine-protease granzymes (Peters et al., 1991). Upon target recognition, LGs are enriched at the IS and are eventually released into the synaptic cleft to induce destruction of target cells (Qu et al., 2011; Stinchcombe et al., 2001). In order to locate their targets, CD8^+^ T cells need to migrate through peripheral tissues featuring a three-dimensional (3D) environment.

Extracellular matrix (ECM) serves as one of the key components in peripheral tissues to scaffold and maintain 3D structures in organs and tissues (Frantz et al., 2010). ECM is a 3D network composed of many fibrous structural proteins, a major constituent of which are collagen proteins, the most abundant protein in mammals (Yue, 2014). A rich body of compelling evidence shows that ECM has a profound impact on cell proliferation, differentiation and tumorigenesis (Daley et al., 2008; Nallanthighal et al., 2019; Walker et al., 2018). Solid tumors are often composed of a condensed ECM compared to the neighboring tissues. This dense ECM creates a physical hindrance to impair infiltration of immune cells into the tumors (Salmon et al., 2012). In a high-density matrix, proliferation of T cells is reduced, and the infiltration of CD8^+^ T cells into tumors featured with high collagen-density is impaired, accompanied with diminished cytotoxicity (Kuczek et al., 2019; Peng et al., 2020). The cytoskeleton plays a pivotal role in regulating migration of CD8^+^ T cells (Acuto and Cantrell, 2000; Dupre et al., 2015). It is mainly composed of actin filaments (F-actin), the microtubule network and intermediate filaments (Etienne-Manneville, 2004; Hohmann and Dehghani, 2019). CD8^+^ T cells migrate in an amoeba-like behavior. During migration, T cells generate protrusions at the leading edge, which are mainly driven by new formation of F-actin (Krummel et al., 2016). The contraction of F-actin mediated by myosin generates force to retract the uropod, the rear part of the cell, enabling CD8^+^ T cells to move forward (Jacobelli et al., 2010). Myosin is an actin-associated motor protein, forming bipolar filaments to bind two actin filaments and enable counter movement of these two actin filaments at the expense of ATP (Sellers, 2000; Sweeney and Houdusse, 2010). In T cells, myosin IIA is the predominantly expressed member of the myosin family (Krummel et al., 2016). Blockage of myosin IIA activity results in deficiency of uropod retraction and therefore significantly impairs T cell migration (Jacobelli et al., 2010). During T cell migration, the microtubule-organizing center (MTOC) is located at the uropod (Ratner et al., 1997). Abrogation of microtubule polymerization does not hinder T cell migration (Billadeau et al., 2007).

Nuclear deformability serves as a limiting factor of cell migration through physical restricted space in 3D (Renkawitz et al., 2019; Wolf et al., 2013). To pass the restricted space between pillars, or narrow channel, the nucleus has to be deformed by force from cytoskeleton to fit the size of space (Denais et al., 2016; Thiam et al., 2016; Wolf et al., 2013), which also facilitates cell migration along the path of least resistance in a complex environment (Renkawitz et al., 2019). Further reports showed that the nucleus shape and its shape changes are correlated with the velocity of cell migration (Kim et al., 2014; Krause et al., 2019).

In this work, we used different densities of ECM to mimic physiological and pathological microenvironments to investigate the impact of dense ECM on CTL-mediated cytotoxicity against tumor cells and to understand the underlying mechanisms. We found that CTL cytotoxicity is substantially reduced in dense collagen matrices. The killing machinery *per se* remains intact whereas migration of CTLs is significantly impaired. Furthermore, perturbation of either actin or myosin IIA led to significantly reduced velocity and persistence, indicating that actomyosin contraction is the driving force for 3D CTL migration. By contrast, disruption of the microtubule network promoted 3D CTL migration. Interestingly, CTLs migrating in dense ECM exhibit deformed nuclei, the extent of which is correlated with migration velocity, indicating that flexibility of CTL nuclei is pivotal to CTL migration in 3D. We found that nuclei flexibility is regulated by the microtubule networks. Finally, our results show that disruption of microtubule but not actin polymerization can rescue the impaired migration as well as the reduced cytotoxic efficiency of CTLs in dense collagen matrices.

## Materials and Methods

### Antibodies and reagents

All chemicals not specifically mentioned are from Sigma-Aldrich (highest grade). All inhibitors not specifically mentioned are from Cayman Chemical. The following antibodies were used: Alexa Fluor 647 anti-human CD3 antibody (UCHT1, BioLegend), Alexa Fluor 488 anti-human Granzyme A antibody (CB9, BioLegend), Alexa Fluor 647 or Brilliant Violet (BV) 510 anti-human perforin antibody (dG9, BioLegend), Alexa Fluor 647 anti-human Granzyme B antibody (GB11, BioLegend), Brilliant Violet (BV) 421 anti-human CD178 (Fas-L), anti α-Tubulin mAb antibody (DM1A, Cell Signaling Technology), and Alexa Fluor 405 conjugated Goat anti-Mouse IgG (H+L) cross-absorbed secondary antibody (ThermoFisher Scientific). All isotype controls of fluorescence conjugated primary antibodies are from BioLegend. The following reagents were used: Alexa Fluor 488 or Alexa Fluor 568 Phalloidin (ThermoFisher Scientific), Atto 488 NHS ester ((ThermoFisher Scientific), Collagenase Type I (ThermoFisher Scientific), FibriCol^®^ Type I Collagen Solution (Bovine, Advanced Biomatrix).

### DNA constructs

For pGK-puro-pCasper-pMax (referred to as pCasper-pMax in the manuscript), the vector backbone used for generation of this plasmid is a kind gift from Ulrich Wissenbach (Saarland University) who previously modified the AMAXA vector (Lonza) by replacing the sequence encoding GFP with a linker sequence encoding a multiple cloning site (pMAX). In a first step, the sequence encoding pCasper was amplified from pCasper3-GR (evrogen #FP971) with the following primers introducing an XhoI recognition site at both ends of the amplicon. Forward primer: 5‘-CTCGAGGCCACCATGGTGAGCGAG-3‘, reverse primer: rev 5‘-GACGAGCTGTACCGCTGACTCGAG-3‘. The amplicon was subcloned into the XhoI site of pMAX. In a second step, the sequence encoding puromycin resistance was introduced into the intermediate plasmid under the control of 3-phosphoglycerate kinase promoter (PGK-1).

The PGK-1-Puromycin sequence was amplified out of pGK-Puro-MO70 vector backbone (Alansary et al., BBA 2015) using the following primers introducing SacI recognition sites at both ends of the amplicon to insert it into the Sac-I site of the intermediate plasmid. Forward primer: 5’-GAGCTCAATTCTACCGGGTAGGGGA-3’, reverse primer 5’-GCAAGCCCGGTGCCTGAGAGCTC-3’. The final plasmid is named pGK-puro-pCasper-pMAX. Final and intermediate plasmids were controlled by endonuclease digestion patterns and sequencing. EMTP-3×GFP was a gift from William Bement (Addgene plasmid # 26741). Histone 2B-GFP was a gift from Geoff Wahl (Addgene plasmid # 11680). LifeAct-mRuby was a kind gift from Roland Wedlich-Söldner (University of Muenster).

### CTL preparation, cell culture, and nucleofection

Peripheral blood mononuclear cells (PBMCs) were obtained from healthy donors as described before (Kummerow et al., 2014). Human primary CD8^+^ T Cells were negatively isolated from PBMCs using Dynabeads™ Untouched™ Human CD8 T Cells Kit (ThermoFisher Scientific) or Human CD8^+^ T Cell Isolation Kit (Miltenyi Biotec), stimulated with Dynabeads™ Human T-Activator CD3/CD28 (ThermoFisher Scientific) with 17 ng/ml of recombinant human IL-2 (ThermoFisher Scientific). MART-1-specific CD8^+^ T-cell clones were generated by Friedmann et al (Friedmann et al., 2020). All CD8^+^ T cells were cultured in AIM V medium (ThermoFisher Scientific) containing 10% FCS and 1% Penicillin-Streptomycin. For nucleofection, CD3/CD28 beads were removed 48 hours after stimulation and 5 × 10^6^ CTLs were electroporated with 2 μg plasmid using 4D-Nucleofector (Lonza). Medium was changed 6 hours after nucleofection and transfected cells were used 24-36 hours after electroporation. Raji and NALM-6 cells were cultured in RPMI-1640 medium (ThermoFisher Scientific) containing 10% FCS and 1% Penicillin-Streptomycin. NALM-6 pCasper cells were generated by Knörck et al (Knörck et al., 2021) and were cultured in RPMI-1640 in the presence of puromycin (0.2 µg/ml). SK-MEL-5 cells were transfected with pCasper-pMax using jetOPTIMUS^®^ DNA Transfection Reagent (Polyplus-transfection) following the manufacturer’s instructions and then cultured in MEM medium (ThermoFisher Scientific) containing 10% FCS and 1% penicillin-streptomycin. All cells were cultured at 37°C with 5% CO_2_.

### Preparation of collagen matrix and cell embedding

As described previously (Schoppmeyer et al., 2018), collagen type I stock solution (10 mg/ml, bovine if not mentioned otherwise) was neutralized with 0.1 N NaOH solution on ice to reach pH 7.0-7.4). Then, 10×PBS was added into the neutralized collagen with a dilution factor of 1:10. This collagen solution was further diluted with PBS to the desired concentrations. Cells of interest were then resuspended in the corresponding collagen solution and collagen was polymerized at 37°C with 5% CO_2_ for 1 hour if not mentioned otherwise. To increase collagen stiffness with ribose, as described by Mason et al (Mason et al., 2013), collagen stock solution was first diluted to a concentration of 3 mg/ml with 0.1% acetic acid with or without 100 mM ribose. These collagen solutions were kept at 4°C for 5 days followed by the normal collagen preparation procedure as described above.

### Killing assay in 3D with the high-content imaging setup

Cells of interest were resuspended in freshly-neutralized chilled collagen solution with indicated concentration. 20 μl of this cell/collagen mixture was added in one well of pre-cooled Corning™ 96-Well half-area plate (Merck) and the plate was centrifuged. The plate was kept at 37°C with 5% CO_2_ for 1 hour for collagen polymerization. Afterwards, 100 µl of AIMV (10% FCS) medium was added to each well, which could contain inhibitors or vehicle control as indicated in the figure legends.

For killing assays, we used either NALM-6 cells stably expressing apoptosis reporter pCasper-pMax or SK-MEL-5 cells transiently transfected with pCasper-pMax (referred to as NALM-6-pCasper or SK-Mel-5-pCasper, respectively) as target cells. NALM-6-pCasper were pulsed with staphylococcal enterotoxin A (SEA, 0.1 µg/ml) and SEB (0.1 µg/ml) at 37°C with 5% CO_2_ for 40 min prior to killing assays. Images were acquired by ImageXpress (Molecular Devices) with Spectra X LED illumination (Lumencor) at 37°C with 5% CO_2_ for 12 to 24 hours with an interval of 15 or 30 min. As described previously (Backes et al., 2018), fluorescence of pCasper-pMax was acquired using LEDs 470/24 for excitation and the following filter sets (Semrock): Ex 472/30 nm, Em 520/35 nm for GFP and Em 641/75 nm for RFP/FRET. A 20× S Fluor 0.75 numerical aperture objective (Nikon) was used. The killing efficiency was calculated as (1-N_exp_(t)/(N_exp_(t_0_)×N_live_(t)/N_live_(t_0_))×100%.

(N_live_: number of live target cells in the control wells without CTLs; N_exp_: number of live target cells in the experimental wells; t_0_: the first time point of the measurement; t: end time point of the measurement).

### Immunostaining and flow cytometry

Cells were fixed with pre-chilled 4% PFA and then washed twice with PBS/0.5% BSA. Then cells were permeabilized and blocked with 0.1% saponin in PBS containing 5% FCS and 0.5% BSA, and then stained with the indicated primary antibody or Alexa Fluor 488 Phalloidin for 30 min at room temperature followed by staining of Alexa Fluor 405 labeled secondary antibody if the primary antibody was not fluorophore-conjugated. Flow cytometry data were acquired using a FACSVerse™ flow cytometer (BD Biosciences) and were analyzed with FlowJo v10 (FLOWJO, LLC).

### Live-cell 3D imaging using light-sheet microscopy

As described previously (Schoppmeyer et al., 2018), collagen with 10 × 10^6^ CTLs/ml polymerized in the capillary at 37°C with 5% CO_2_ for 2 hours. Afterwards, the samples were scanned with light-sheet microscopy Z1 (Zeiss) at 37°C for 30 min with an interval of 30 sec and a z-step size of 1 µm. A 20× objective (W Plan-Apochromat, N.A. 1.0) was used. Excitation was realized by two lasers, 488 and 561 nm. Emission was filtered via Em525/40nm and Em 585 LP filters. The images were acquired with ZEN software. Trajectories of CTLs and nuclear irregularity index (NII) were tracked and analyzed with Imaris 8.1.2 (containing Imaris, ImarisTrack, ImarisMeasurementPro, ImarisVantage from Bitplane AG). The nuclei or the cell bodies were detected automatically by ImageJ/Fiji based on the corresponding fluorescence, and parameters (circularity and Feret’s diameter) were analyzed with ImageJ/Fiji. *Visualization of CTL migration in a planar 3D collagen matrix with Zeiss Observer Z.1* CTLs (5 × 10^6^ cells/ml) were resuspended in collagen solution with or without Calcein labeled target cells (5 × 10^6^ cells/ml). Cell/collagen mixture (3 µl) was pipetted as a droplet onto the center of an Ibidi μ-dish (Ibidi GmbH). Then a Sigmacote^®^ (Merck) coated glass coverslip (5 mm, Orsatec GmbH) was carefully placed on top to flatten the droplet (calculated thickness around 150 µm). The Ibidi μ-dish was closed with the lid and incubated at 37°C with 5% CO_2_ for 1 hour. After collagen polymerization, the glass coverslip was removed from collagen matrix. For migration assay, CTLs were either non-fluorescent (in presence with target cells) or stained with 1 µg/ml Hoechst 33342 in AIMV (10% FCS) medium for 30 min (without target cells), and then were incubated in fresh AIMV (10% FCS) medium for another 30 min in presence of inhibitors or vehicles as indicated in the figure legends. Raji cells and unlabeled CTLs were mixed The images were acquired with Zeiss Observer Z.1 either every 30 sec for 30 min or every 1 min for 3 hours at 37°C with 5% CO_2_ with a Zeiss Colibri LED illumination system and 10× objectives (Fluar 10× /0.25 M27 Air). Images were taken using an AxioCamM1 CCD camera and AxioVision 4.1.8. The images of cell migration and nuclear irregularity index (NII, measured as nuclear circularity) were analyzed using Imaris 8.1.2. Migration trajectories in presence of target cells were analyzed by Fiji. To quantify CTL search efficiency, CTLs were randomly selected from the CTLs that were observed for the whole period. The probability for CTLs finding at least one target within 3 hours was quantified. From these CTLs, the time of CTL contacting the first target cells was quantified, if this CTL could find at least one target cell within 3 hours.

### Confocal microscopy

For the fixed sample, CTLs were fixed at indicated time points with 4% PFA and permeabilized with 0.3% Triton-100 with 5% FCS in PBS, followed by staining with indicated antibodies or fluorescent dyes according to the manufacturers’ instructions. Images were acquired by confocal microscopy LSM 710 with a 63× objective (N.A. 1.4) and a Nikon E600 camera using ZEN software. The nuclear irregularity index (NII, measured as nuclear circularity from maximum intensity projection) was analyzed with ImageJ.

### Visualization of collagen structure and determination of porosity

After collagen polymerization in the capillary, collagen was stained with Atto 488 NHS ester in PBS (50 µM) at room temperature for 15 min. Afterwards, the collagen matrix was washed by PBS twice. Matrix structure of collagen was visualized by light-sheet microscopy with a 20× objective (W Plan-Apochromat, N.A. 1.0). Collagen pore size was measured in the middle slice of the z-stack by Fiji (BIOP version) with Max Inscribed Circles plugin as described elsewhere (Acton et al., 2014).

### Shear Rheology Stiffness measurement

Rheology measurements with different concentrations of bovine collagen were performed using DHRIII Rheometer (TA Instruments). For figure 1.b, 50 µl of the neutralized collagen solution (pH 7.0-7.4) was placed between two parallel plates of 12 mm diameter pre-heated to either 25°C or 37°C to polymerized collagen solutions. The shear moduli were measured at frequency ω – 3 rad/s while the temperature was at 37°C as described previously (Madsen and Cox, 2017). All experiments were performed in triplicates.

**Figure 1.**
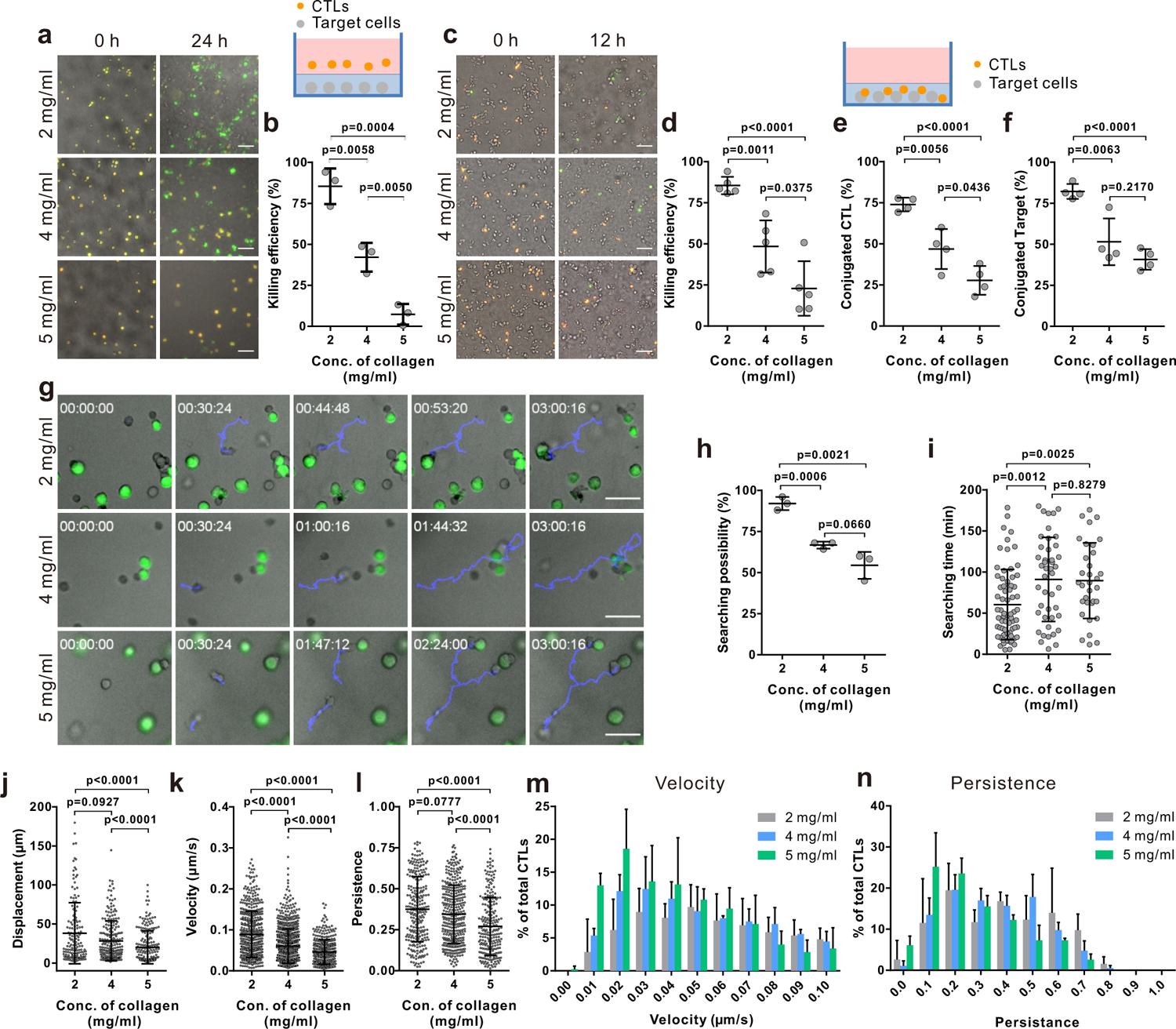
The killing efficiency of CTLs is substantially decreased in dense ECM. **a-d** CTL-mediated cytotoxicity is drastically impaired in dense ECM. Target cells (SEA/SEB pulsed NAML-6-pCasper) were embedded in collagen in absence (**a, b,** 3 donors) or presence of CTLs (**c, d,** 5 donors). Images were acquired using ImageXpress with Spectra X LED illumination (Lumencor) for 24 (**a, b**) or 12 hours (**c, d**). Orange and green color indicates live and apoptotic target cells, respectively. **e-f** Fraction of CTLs in conjugation with target cells (**e**) or target cells in conjugation with CTLs (**f**) was analyzed at 1 hour after polymerization (4 donors). **g-i** Searching efficiency of CTLs is reduced in dense ECM. Target cells (SEA/SEB pulsed Raji) were loaded with calcein (green) and embedded in planar collagen with CTLs. Migration was visualized using Cell Observer (10× objective) at 37°C with an interval of 30 sec for 3 hours. CTLs were tracked manually and representative tracks are shown in **g** (blue lines). and the likelihood to find a target within 3 hours from **g** are shown in **h** (3 donors). The cells finding at least one target cell from **h** were measured for the time required finding their first target (**i**). **j-n** Characterization of CTL behavior in 3D collagen. CTLs were transfected with histone 2B-GFP and LifeAct-mRuby, and then embedded in 3D collagen. Migration was visualized using light-sheet microscopy (20× objective) for 30 min with an interval of 30 sec. The nuclei position were tracked with Imaris to determine the displacement, velocity, and persistence of CTLs shown in **j**, **k**, and **l**, respectively (3 donors). The distribution of CTL velocity and persistence is shown in **m** and **n**, respectively. One dot represents one donor (in **b, d, e, f**, and **h**) or one cell (in **i-l** from 3 donors). Results are presented as Mean±SD. P values were assessed using P values in **i-l** were assessed using the Mann-Whitney test. Other P values were assessed using the unpaired Student’s t-test. Scale bars are 50 μm.

### Statistical analysis

Data are presented as mean ± SD. GraphPad Prism 6 Software (San Diego, CA, USA) was used for statistical analysis. The data were tested whether they fitted Gaussian distribution. If data fit Gaussian distribution, the differences between two columns were analyzed by the Student’s t-test. If data do not fit Gaussian distribution, the differences between two columns were analyzed by the Mann-Whitney test. If the number of data is too small to test Gaussian distribution, the differences between two columns were analyzed by the Student’s t-test.

### Online Supplemental material

Supplemental Figures 1-4, Video 1-7 and the corresponding legends can be found in Supplemental material. Supplemental Figure 1 shows characterization of target lysis in 3D by CTLs. Supplemental Figure 2 shows impact of collagen density on lytic granule pathway. Supplemental Figure 3 shows structure of type I collagen. Supplemental Figure 4 shows FAK and ROCK are essential for CTL migration in 3D. Movie 1-3 show CTL migration trajectories measured by light-sheet microscopy in 2 mg/ml (Movie 1), 4 mg/ml (Movie 2), and 5 mg/ml (Movie 3) collagen matrices. Movie 4-6 show Nuclei shape dynamics of CTLs during migration in 2 mg/ml (Movie 4), 4 mg/ml (Movie 5), or 5 mg/ml collagen (Movie 6). Movie 7 shows microtubule network (green) and F-actin dynamic (red) during CTL migration in 4 mg/ml collagen matrices.

## Results

### CTL cytotoxicity and motility is substantially impaired in dense collagen

To investigate the impact of collagen density on the killing efficiency of CTLs, we used bovine collagen at three concentrations (2 mg/ml, 4 mg/ml, and 5 mg/ml). Primary human CD8^+^ T cells were stimulated with anti-CD3/anti-CD28 antibody-coated beads to obtain CTLs. The target cells were embedded in collagen matrices and CTLs were settled on top of the gels as depicted in Fig. 1a. Target cells ((NALM-6-pCasper) stably express apoptosis reporter pCasper-pMax, a GFP-RFP FRET pair linked by a consequence containing caspase recognition site (DEVD), allowing the detection of cell death. As reported previously (Backes et al., 2018), CTL can induce target cell apoptosis (cleavage of the linker between the GFP-RFP FRET pair leading to a switch of fluorescence to green) or target cell necrosis (loss of fluorescence) (Supplemental Fig. 1a). Using a high-content imaging setup (ImageXpress), we observed that after 24 hours in 2 mg/ml collagen, 85.4±10.9% of target cells were killed by either apoptosis or necrosis. In 4 mg/ml collagen, the fraction of apoptotic and necrotic target cells combined is down to 42.1±8.8%; remarkably, in 5 mg/ml collagen, only 7.3±6.3% was eliminated by CTLs (Fig. 1a, 1b). The same tendency holds true for earlier time point 12 hours (example in Supplemental Fig. 1b). These results indicate that CTL killing efficiency was substantially diminished in dense collagen.

In the above-mentioned setting, CTLs had to infiltrate into the matrix. To exclude a possible influence of the initial infiltration phase on cytotoxic efficiency, we embedded CTLs with target cells in the collagen matrix. The cytotoxic efficiency of CTLs was again different for the different collagen densities (Fig. 1c, 1d) and was very similar to the conditions which included infiltration (Fig. 1a, 1b). Therefore, the cytotoxic efficiency of CTLs is strongly reduced in dense collagen and this effect is independent of matrix infiltration.

To unravel potential mechanisms of reduced CTL cytotoxicity in dense collagen, we tested different possibilities. First, the expression of cytotoxic proteins involved in CTL cytotoxicity was tested. We harvested CTLs from collagen matrices using collagenase. No significant differences in the expression of cytotoxic proteins (perforin, granzyme A and granzyme B) were observed for the three different concentrations (Supplemental Fig. 2a). FasL expression was also examined, which was very low for all three concentrations (Supplemental Fig. 2b). In addition, control experiments show that the short-time treatment of collagenase does not alter the protein level on T cells, e.g. CD3 expression (Supplemental Fig. 2c). Interestingly, in dense collagen, the fraction of CTLs conjugated with targets (Fig. 1e) and the fraction of target cells conjugated with CTLs (Fig. 1f) were significantly reduced. These results indicate that the impaired CTL killing efficiency in dense collagen is not owed to any change in the main components of the killing machinery but rather to reduced search efficiency.

**Figure 2.**
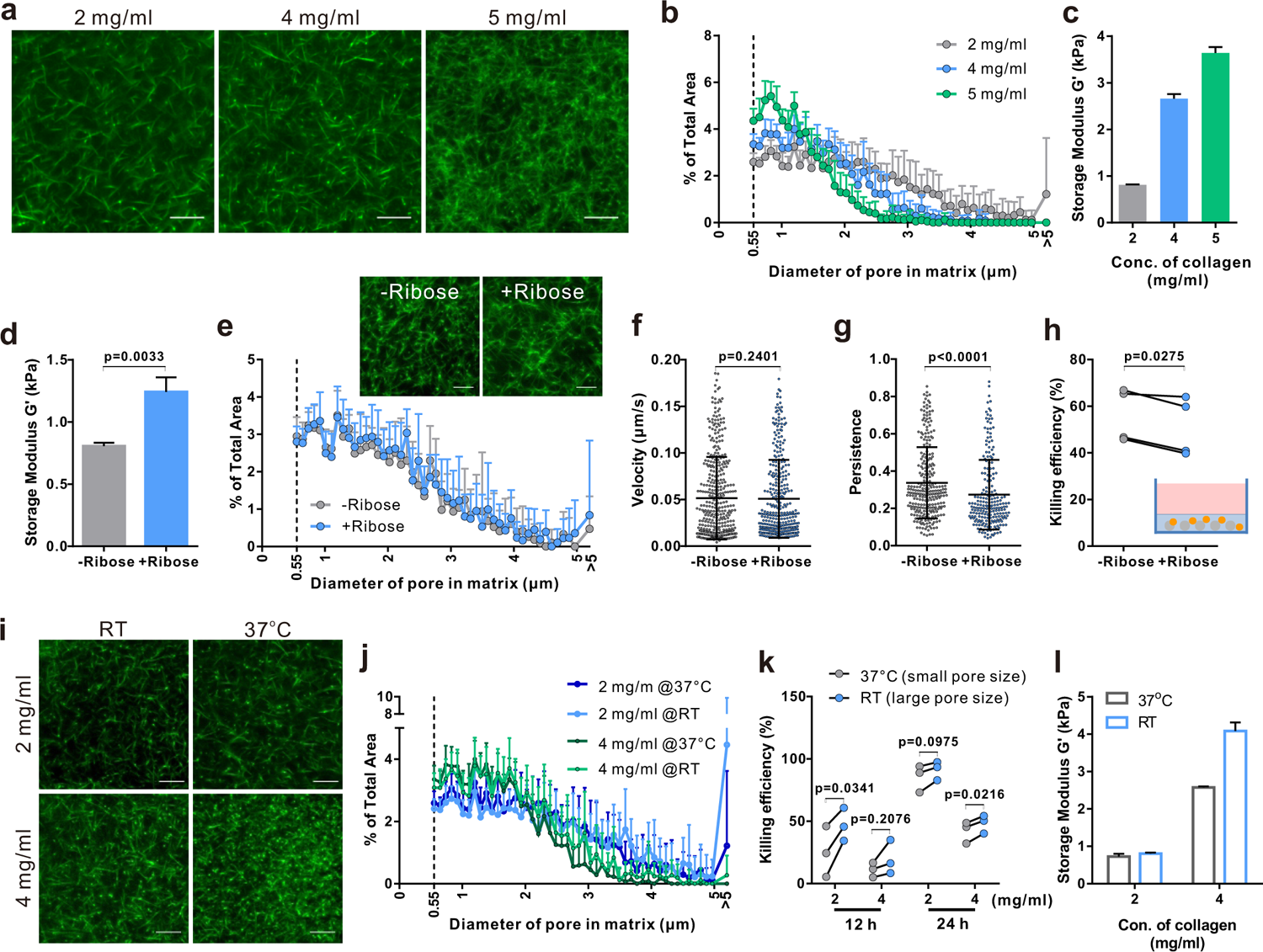
Both enhanced stiffness and reduced pore size contribute to impaired CTL killing efficiency in dense ECM. **a-b** Characterization of porosity in collagen matrices with different densities. Type I bovine collagen was stained with Atto 488 NHS ester, and visualized using light-sheet microscopy with a 20× objective (**a**). The pore size from **a** was analyzed and the distribution is shown in **b**. **c** Stiffness of collagen matrices with different densities. The storage modulus of the respective collagen matrix was measured at frequency ω – 3 rad/s (37°C). Results are mean±SD from three independent experiments. **d-h** Adding ribose increases stiffness of collagen (2 mg/ml) without affecting the pore size. Ribose or vehicle control was added to collagen for five days prior to collagen polymerization (2 mg/ml). Stiffness and pore size of the respective conditions is shown in **d** and **e**, respectively. **f-h** Stiffening collagen leads to reduced CTL killing efficiency in 3D. Hoechst 33342-stained CTLs were embedded in a planar 3D collagen matrix. Migration was visualized using Cell Observer (10× objective) for 30 min with an interval of 30 sec (**f**, **g**). Migration velocity and persistence are shown in **f** and **g**, respectively. To determine killing efficiency, CTLs were embedded with NALM6-pCasper. Killing efficiency was determined after 4 hours and is shown in **h**. **i-j** Pore size is increased by polymerizing collagen at RT. Bovine type I collagen (2 mg/ml or 4 mg/ml) stained with Atto 488 NHS ester was either polymerized at 37°C for 1 hour or at RT for 30 min and then at 37°C for 30 min. The structure was visualized using light-sheet microscopy (20× objective) (**i**). The pore size at each condition is determined and shown in **j**. **k** Enlarged pore size improves CTL killing in 3D. NALM6-pCasper cells were embedded at the indicated conditions and CTLs were added from the top. **l** Characterization of stiffness of collagen polymerized at different temperatures. The stiffness was determined as in **d**. One dot stands for one cell (in **f** and **g**) or one donor (in **h** and **k**). Scale bars are 10 μm. Results of **f, g, h, k** are from 3 donors. Other results are from 3 independent experiments. Results presented as Mean±SD. P values in **d** were assessed using the unpaired Student’s t-test. P values in **f**, **g** were assessed using the Mann-Whitney test. P values in **h**, **k** were assessed using the paired Student’s t-test.

We next investigated the search efficiency of CTLs in collagen in detail using live-cell imaging. The example in Fig. 1g shows that the CTL highlighted by the blue track in a 2 mg/ml collagen needs around 45 min to find the first target, whereas the CTLs in 4 mg/ml and 5 mg/ml ECM need about 100 min (lower panels of Fig. 1g). Notably, in collagen with low density (2 mg/ml), all most all CTLs (92.0±4.0%) found at least one target cell within 3 hours; whereas in dense collagen this likelihood (66.8±2.1% or 54.4±8.2%) reduced with the increased density (4 mg/ml and 5 mg/ml, respectively, Fig. 1h). In comparison, for the CTLs that found target cells, to find the first target, CTLs needed about 90 min in 4 mg/ml and 5 mg/ml collagen but only about 60 min in 2 mg/ml collagen (Fig.1i). These results suggest that the possibility for CTLs to find their targets is decreased in dense collagen.

Appropriate migration of CTLs is one key factor for optimal search efficiency. We, therefore, examined CTL motility in collagen matrices. To precisely quantify migration parameters we first analyzed displacement, velocity, and persistence. Using light-sheet microscopy, we observed that in dense collagen matrices (4 mg/ml and 5 mg/ml), the displacement of CTLs (distance between the starting point and the end point) was reduced (Fig. 1j, Movie 1-3). In addition, analysis of trajectories in 3D collagen shows that both velocity and persistence of CTL migration were decreased in a concentration-dependent manner (Fig. 1k, l,). Analysis of velocity and persistence distributions reveals that the fraction of CTLs with low velocity and low persistence is higher in dense ECM (5 mg/ml, Fig. 1m, n). In summary, our findings suggest that in a dense matrix, CTL migration is hindered, which is likely the reason for longer search time and reduced cytotoxic efficiency.

### CTL migration in 3D is regulated by collagen stiffness and porosity

To examine the underlying mechanisms, we turned to the physical properties of collagen matrix including stiffness and porosity (pore size). We fluorescently labeled collagen to visualize its structure (Fig. 2a). Pores between collagen fibers were observed and measured (Supplemental Fig. 3), the size of which was decreased with increasing density (Fig. 2a, b). Collagen stiffness was determined with rheology measurements, which is increased with collagen density (Fig. 2c). This observation of smaller pore size and higher stiffness in dense collagen is in good agreement with the others (Lang et al., 2015; Wolf et al., 2013). To study which of these two features contributes to the impaired CTL migration in collagen matrix, we modified the collagen matrix. In terms of the stiffness, the addition of 100 mM ribose has been reported to increase the stiffness of rat collagen without changing the concentration (Mason et al., 2013). We reproduced these conditions with bovine collagen and found the same tendency: with the addition of 100 mM ribose, the collagen stiffness was increased as indicated by the storage modulus (Fig. 2d), without affecting the pore sizes (Fig. 2e). Upon increase in stiffness of the collagen, CTL velocity was not altered (Fig. 2f), whereas the persistence was reduced (Fig. 2g). In line with this result, the cytotoxic efficiency of CTLs was reduced in the presence of ribose (Fig. 2h). These results indicate that stiffness is involved in regulating CTL migration persistence, which could influence the correlated CTL cytotoxicity.

**Figure 3.**
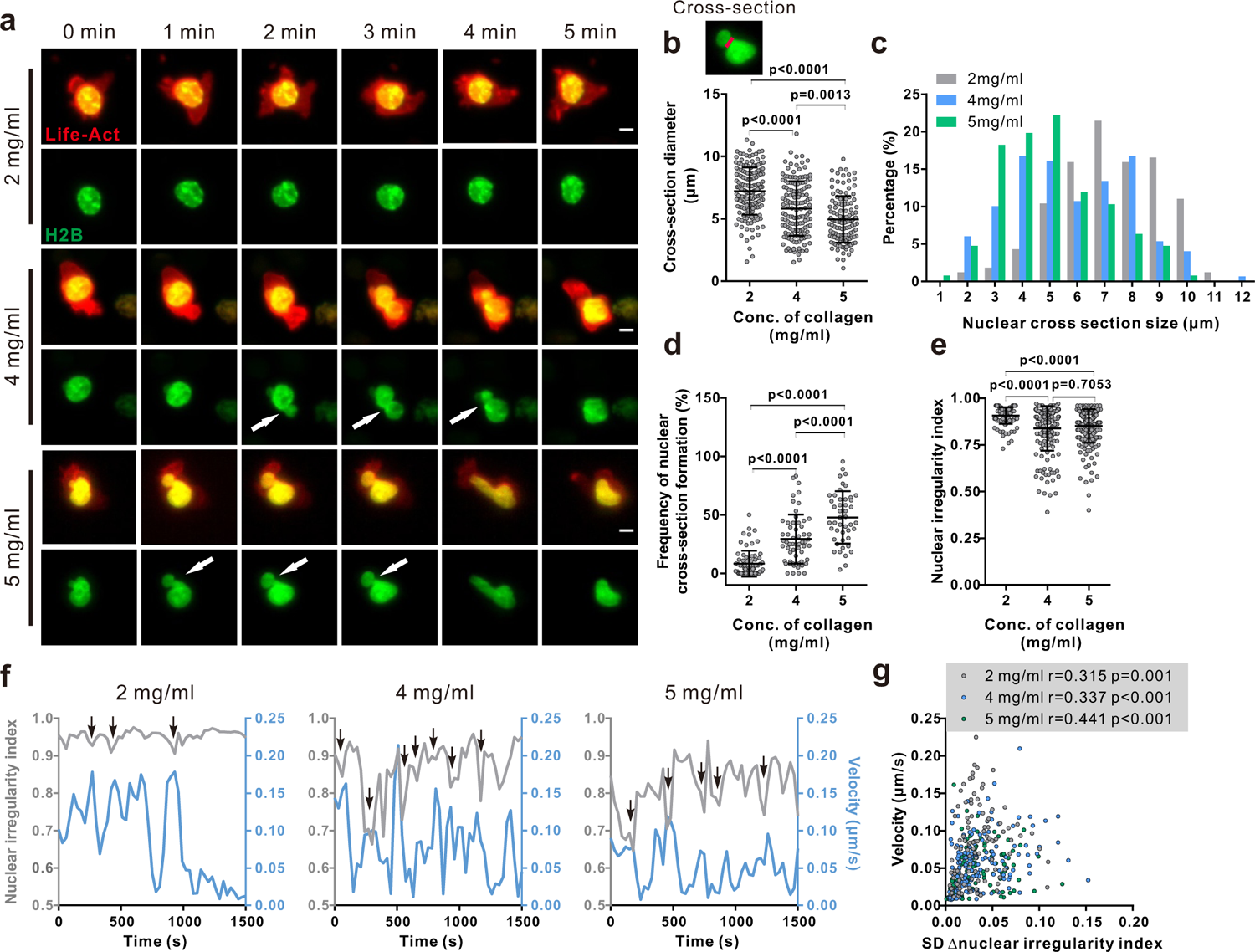
Nuclear deformation is correlated with migration velocity in dense ECM. **a-b** Nuclear deformation in CTLs is enhanced upon the increase in ECM density. CTLs transfected with Histone 2B-GFP (green) and LifeAct-mRuby (red) were embedded in collagen. Migration was visualized using light-sheet microscopy (20× objective) at 37°C for 30 min with an interval of 30 sec. Exemplary cells are shown in **a**. Cross-sections are pointed by the arrowheads. Scale bars are 5 µm. Quantification of cross-section diameter (indicated as red-line in the inset) is shown in **b**. Comparison of distribution of cross-sections diameter in collagen with different densities and the duration of cross-sections are shown in **c** and **d**, respectively. **e** The nuclear irregularity index (sphericity) was reduced with the density of collagen. The nuclear irregularity index from **a** was analyzed. **f** Migration velocity of CTLs is positively correlated with nuclear deformation. Nuclear deformation is determined by the nuclear irregularity index (sphericity). **g** Average velocity is positively correlated with the mean of the change of the nuclear irregularity index. One dot represents one cell. Correlation coefficient is analyzed with Spearman’s correlation. Results are from 3 donors and presented as Mean±SD for **b**, **d**, and **e**. P values were assessed using the Mann-Whitney test.

To change the pore size of a matrix at identical collagen concentrations, we followed a protocol shown to enlarge pore sizes by polymerizing the collagen matrix at room temperature instead of 37°C (Shannon et al., 2015). The pore sizes at two different collagen concentrations (2 mg/ml and 4 mg/ml) polymerized at room temperature or 37°C were compared. We found that the fraction of big pores (diameter larger than 3 µm) was increased at both concentrations, especially at 4 mg/ml (Fig. 2i, 2j). Remarkably, when the pore sizes were enlarged, the cytotoxic efficiency of CTL was enhanced for both concentrations (Fig. 2k). Noticeably, with enlarged pores, the stiffness was not changed for collagen with low density (2 mg/ml) but was increased for dense collagen (4 mg/ml) (Fig. 2l), which could be attributed to better fibrillation kinetics of collagen at RT (Leikina et al., 2002; Zhu and Kaufman, 2014). It indicates that the contribution of enlarged pore size *per se* in CTL killing in the dense matrix could be underestimated due to the increased stiffness. Together, these results suggest that both the pore size and the stiffness are responsible for impaired CTL cytotoxicity resulting from hindered migration in dense ECM, whereby CTL migration persistence is mainly determined by matrix stiffness while the velocity is likely determined by the pore size of the matrix.

### Deformability of nucleus is a limiting factor for CTL migration in dense ECM

As the stiffest organelle in cells, the nucleus is essential for decision making of migration direction for immune cells migrating through restricted space (Denais et al., 2016; Thiam et al., 2016; Wolf et al., 2013). Therefore, we examined the role of the nucleus in impaired CTL migration in dense ECM. First, we noticed that the nuclear morphology was deformed to an hour-glass shape in migrating CTLs in ECM as shown in the time-lapse (Fig. 3a, Movie 4-6). To quantify the extent of this nuclear deformation, we analyzed the diameter of cross-sections (the shortest intersection of the hour-glass shape), as well as the sphericity (how closely an object resembles a sphere). The cross-section diameter was decreased with increasing collagen density (Fig. 3b, c). Furthermore, in dense collagen, the nuclei of migrating CTLs were more frequently deformed to this hour-glass shape than their counterparts in collagen with low density (Fig. 3d). The analysis of nuclear irregularity index (sphericity) shows that nuclear irregularity index was decreased in dense collagen (4 mg/ml and 5 mg/ml) relative to the collagen with low density (2 mg/ml) (Fig. 3e). It is reported that in dendritic cells, the nucleus is drastically deformed when migrating through spatially restricted areas (Renkawitz et al., 2019). Together, our results suggest that the extent of nuclear deformation is increased in dense collagen likely due to its decreased porosity.

We next examined whether nuclear deformation was correlated with CTL migration in 3D. As shown in the exemplary cells, at the time points when the nucleus was deformed, the real-time velocity was high and this phenomenon was observed for all three concentrations (Fig. 3f). The respective analyses show that the migration velocity of CTLs is positively correlated to nuclear deformation as determined by the change in nuclear irregularity index (sphericity) in all three densities (Fig. 3g). This observation is in good agreement with a recent report, showing that cell migration velocity positively correlated to the change in nucleus shape in 2D (Krause et al., 2019). Together, these findings suggest that deformability of the nucleus is a key factor to determine CTL migration in 3D.

### Nuclear deformability is regulated by the microtubule network

Cytoskeleton is an essential regulator for nuclear deformation induced by mechanical forces (Alisafaei et al., 2019; Kim et al., 2015) as well as for cell migration (Serrador et al., 1999). Therefore, we focused on the role of key cytoskeletal components in regulating nuclear deformation and CTL migration in 3D. The expression of F-actin (Fig. 4a) and microtubules (Fig. 4b) in collagen-embedded CTLs was modestly upregulated in the dense matrices (4 mg/ml and 5 mg/ml). We next analyzed the intracellular distribution of F-actin and microtubules using immuno-staining. Confocal images and live-cell imaging show that F-actin was mainly located in the CTL cortex, whereas the microtubule network was enriched around the microtubule-organizing center (MTOC) at the uropod and nucleus-surrounding areas (Fig. 4c, Movie 7) as reported by the others (Kopf et al., 2020). Noticeably, the average distance of microtubule network and the nucleus was decreased upon the increase in collagen density (Fig. 4d, e). As it has been reported that the volume of the nucleus is limited by the microtubule network (Kim et al., 2015), we hypothesized that the microtubule network is involved in regulating CTL nuclear deformability.

To investigate this hypothesis, we used nocodazole, an inhibitor of tubulin polymerization, to abrogate the functionality of the microtubule network. We found that nuclear sphericity was significantly decreased by nocodazole treatment as shown in the exemplary cells (Fig. 4f) and in the quantification of (Fig. 4g), indicating that with microtubules depolymerized, nucleus is more deformable.

Combined with the finding that nuclear deformation is correlated with CTL migration, we postulated that disruption of the microtubule network should impact CTL migration, especially in dense ECM. Analysis of migration of CTLs in collagen matrices shows that in CTLs treated with nocodazole migration velocity was enhanced at all ECM densities (Fig. 4h); whereas persistence was only increased in collagen with 5 mg/ml (Fig. 4i). In comparison, disruption of the actin network by latrunculin-A or abrogation of myosin IIA by blebbstatin almost abolished CTL velocity and persistency for all three collagen densities (Fig. 5h, 5i). Inhibition of a myosin IIA-upstream kinase Rock, or focal adhesion kinase (FAK) also drastically impaired CTL velocity (Supplemental Fig. 4a), but did not drastically change 3D migration persistence (Supplemental Fig. 4b). Furthermore, live-cell imaging shows that the nucleus in the nocodazole-treated CTLs was more deformable than their DMSO-treated counterparts (Fig. 4j). For CTLs, despite nocodazole-or vehicle-treated, the migration velocity was positively correlated with the extent of nuclear deformation (Fig. 4k). In summary, we conclude that disruption of the microtubule network enhances CTL migration especially in dense ECM, which is correlated with enhanced deformability of nucleus in CTLs.

**Figure 4.**
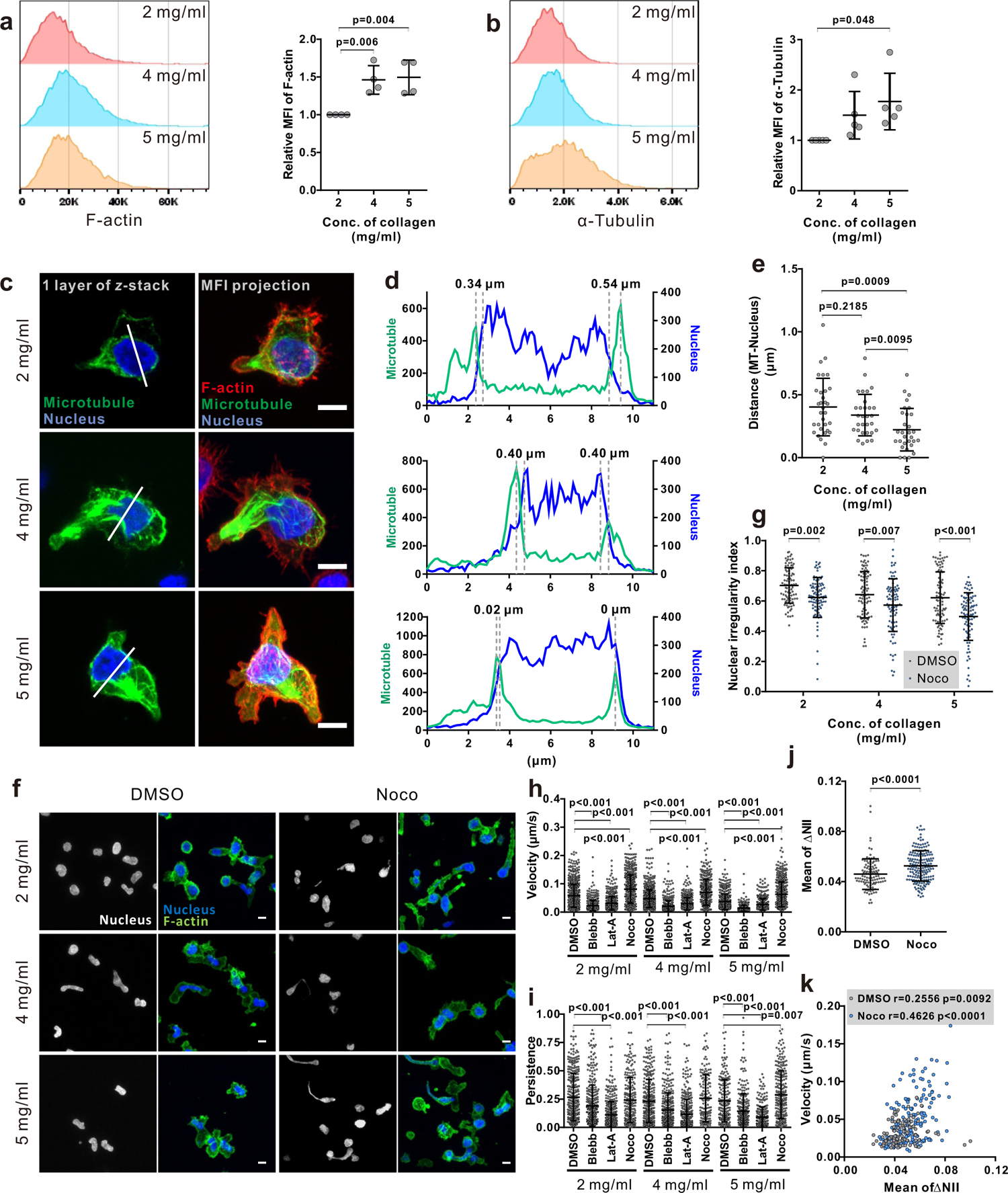
Disruption of microtubule network enhances CTL nuclear deformation and migration in dense ECM. **a-b**, Dense ECM slightly up-regulates expression of F-actin and α-tubulin. CTLs were embedded in the respective collagen matrix for 5 hours after polymerization and then harvested after collagenase digestion, followed by fixation with 4% PFA. The F-actin and α-tubulin were stained with Alexa Fluor 488 Phalloidin and anti α-Tubulin mAb antibody. The samples were measured with flow cytometry. One presentative donor is shown in the left panels and the quantification is shown in the right panels. One dot represents one donor. **c-e** The microtubule network is located at the nucleus-surrounding region. CTLs were transfected with EMTP-3×GFP (green), and 24 hours post-transfection CTLs were fixed and stained with Hoechst 33342 (blue), and Alexa 568-phalloidin (F-actin, red). Images were acquired with confocal microscopy (63× objective). MIP: maximum intensity projection. Scale bars are 5 µm. The fluorescence intensity along the random line cross nucleus depicted in **c** is shown in **d**. Distance between microtubules and nucleus is defined as the distance between two maxima as indicated in **d** and quantified in **e**. **f-g** Disruption of the microtubule network increases the level of nuclear deformation in CTLs. CTLs were embedded in the respective collagen matrix and then treated with nocodazole (Noco, 10 µM) or DMSO for 5 hours prior to fixation. Then CTLs were stained with Hoechst 33342 (blue) and Alexa 488-phalloidin (F-actin, green). Images were acquired with confocal microscopy (63× objective). Exemplary images are shown in **f**. Quantification of the nuclear irregularity index (circularity) in maximum intensity projection is shown in **g**. **h-i** Impact of cytoskeleton components on CTL migration in 3D. Hoechst 33342-stained CTLs were embedded in planar collagen present with DMSO, blebbstatin (Blebb, 50 µM), latrunculin-A (Lat-A, 50 nM), or nocodazole (Noco, 10 µM). Migration was visualized with cell observer (10× objective) at 37°C for 30 min. CTLs were automatically tracked and analyzed with Imaris. Migration velocity and persistence are shown in **h** and **i**, respectively. **j** Disruption of the microtubule network enhances nuclear deformability. CTLs were embedded in collagen (5 mg/ml) and then treated as in **h** and **i**. Nuclear deformability is determined by the change of the nuclear irregularity index (circularity). **k** Correlation of migration velocity and nuclear deformability (the nuclear irregularity index). CTLs from **j** were analyzed. The correlation coefficient is analyzed with Spearman’s correlation. One dot represents one cell. Scale bars are 5 µm. Results are from 3 donors and presented as Mean±SD. P values were assessed using the unpaired Student’s t-test (in **a, b, e**) or the Mann-Whitney test (**g**, **h**, **i**, **j**).

**Figure 5.**
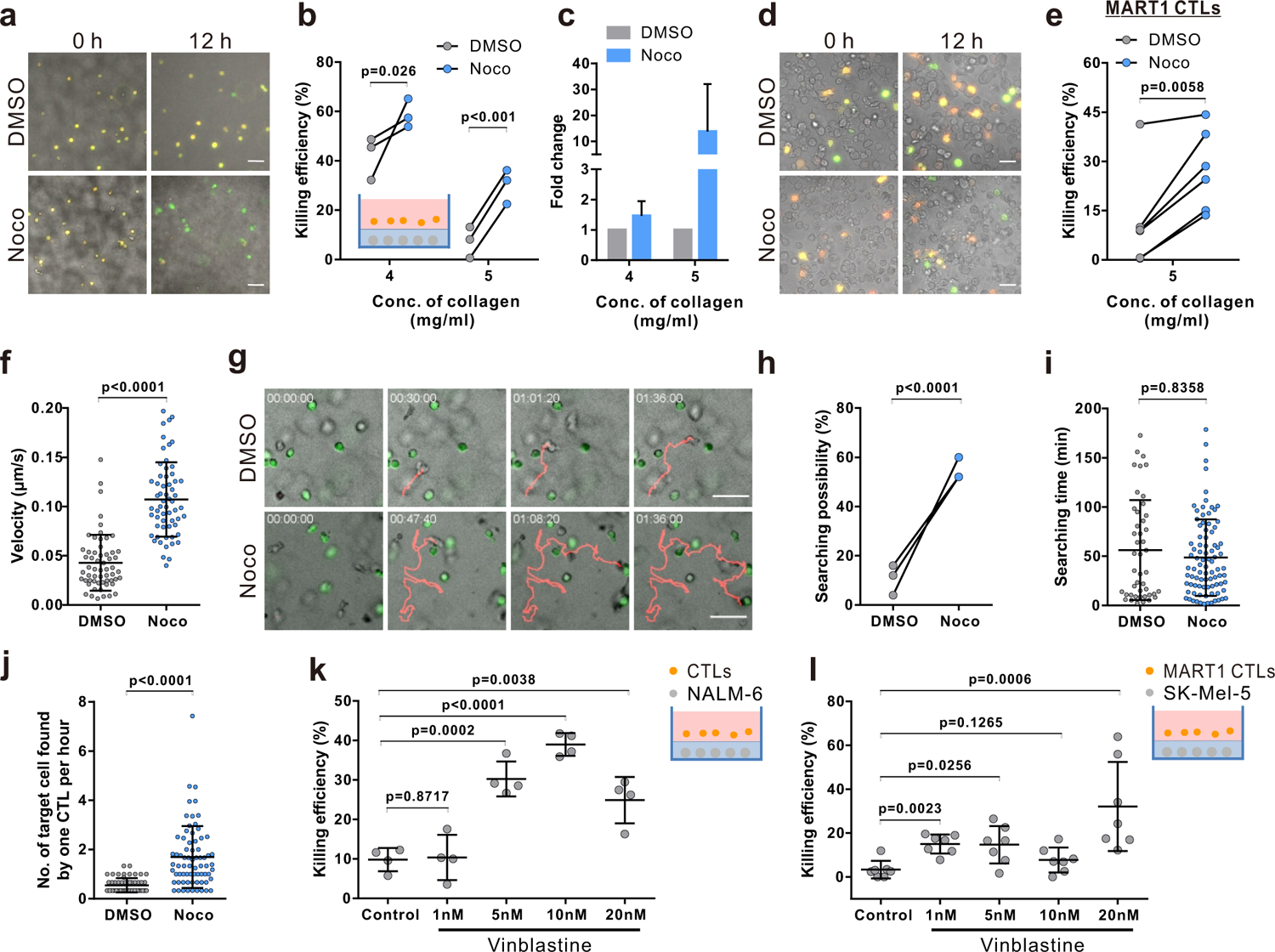
Disruption of microtubules ameliorate impaired CTL killing in dense ECM. **a-c** Inhibition of microtubule polymerization improves dense ECM-impaired CTL killing. Latrunculin-A (Lat-A, 50 nM), nocodazole (Noco, 10 µM), or DMSO were added in the medium with CTLs prior to the killing assay. Images were acquired using Spectra X LED illumination (Lumencor) (20× objective) at 0 and 12 hours. Data are from 3 donors. One representative experiment (5 mg/ml) is shown in **a**. **d-e** Disruption of microtubules improves antigen-specific CTL killing in dense ECM. SK-Mel-5 cells were transiently transfected with pCasper-pMax and were used as target cells. MART1-specific CTLs were added after collagen polymerization as indicated in the inset. One representative experiment (5 mg/ml) is shown in **d**. **f-j** Nocodazole treatment elevates migration and searching efficiency in dense collagen. CTLs were embedded in planer collagen matrix (5 mg/ml) in presence of calcein-loaded target cells (green) and visualized with cell observer. **f** Migration velocity within the duration of 30 min before conjugating with target cells is analyzed. **g** Migration trajectories were tracked manually. The likelihood for CTL to find target cells (**h**) and the time required for CTL to find the first target cell (**i**) are shown. **j** Number of target cells found by one CTL per hour. **k, l** Vinblastin enhances killing efficiency of CTLs in dense ECM. Respective target cells as indicated were embedded in collagen (5 mg/ml). Killing efficiency was determined at 24 hours (**k**) or 12h (**l**) after adding CTLs using ImageXpress (20× objective). One dot represents one donor (in **b**, **h,** and **k**), one dot represents one independent experiment (in **e** and **l**), or one cell (in **f**, **i,** and **j**, from 4 donors). Scale bars are 50 µm. Results are presented as Mean±SD. Scale bars are 50 µm. P values were assessed using the paired Student’s t-test (in **b**, **e**, **h**) or the Mann-Whitney test (in **f** and **i-l**).

### Disruption of microtubules ameliorate impaired CTL killing in dense ECM

Considering the dependence of CTL migration velocity and persistence on the microtubule-network, we hypothesized that interference with microtubules should improve the impaired cytotoxicity in dense ECM. Results of 3D killing assay show that indeed in dense matrix (4 mg/ml and 5 mg/ml), disruption of the microtubule network ameliorated CTL killing (Fig. 5a-c). Moreover, to further confirm this phenomenon, we used primary human CTL clones specific for MART1 (also known as Melan-A). A melanoma cell line SK-MEL-5, which endogenously, present MART1 on their surface (Sugita et al., 1996), was used as target cells. The time lapse shows that in good agreement with previous results, nocodazole-treatment significantly enhanced killing efficiency of CTL clones to remove SK-MEL-5 (Fig. 5d, 5e). Together, our findings suggest that the microtubule network can serve as a promising target to improve CTL killing against tumor cells in dense ECM.

We further confirmed that in presence of target cells, disruption of the microtubule-network also promoted 3D CTL migration (Fig. 5f), as in the case of without target cells (Fig. 4h), Concomitantly, the likelihood for microtubule-disrupted CTLs to find their target cells was drastically enhanced compared to their control counterparts (Fig. 5g, 5h). For those CTLs that did find their targets, nocodazole-treated CTLs spent similar time to locate their first targets as DMSO-treated ones (Fig. 5i). However, on average, the number of targeted cells found by nocodazole-treated CTLs was significantly higher than by control CTLs (Fig. 5j). Finally, we tested vinblastine, a microtubule-inhibitor applied as a chemotherapeutic to disrupt tumor cell mitosis (Tamzali and Kemp-Symonds, 2015). We treated CTLs with vinblastine to determine CTL killing efficiency in dense collagen. We found that vinblastine increases both killing efficiency of primary CTLs (Fig. 5k) and human MART1-specific CTL clones (Fig. 5l). Together, we conclude that disruption of the microtubule network significantly enhances CTL migration and killing efficiency in dense collagen.

## Discussion

Proper motility of CTLs in 3D environments, especially through dense ECM, is the key for search efficacy and the consequent killing efficiency. Two physical properties of ECM are decisive for T cell migration: pore size (porosity) and stiffness (or elasticity) of the fibrils. As concentration of collagen increases, pore size gets smaller and stiffness is elevated. Our work shows that human CTLs migrate spontaneously in 3D collagen matrix. Both the speed and the persistence of their 3D migration diminish along with the increase of collagen concentration. This concentration-dependent correlation of porosity and stiffness can be decoupled. Keeping the concentration of collagen constant, lowering polymerization temperature increases pore size, whereas pretreatment of collagen with ribose enhances the stiffness of collagen fibrils. We show that speed and persistence of human CTL migration in 3D are distinctively regulated: the speed is strongly correlated with the pore size, in good agreement with the observation in murine naive T cells (Hons et al., 2018); while the persistence is mainly determined by the stiffness, in good agreement with a recent report for ovarian cancer cells (Hetmanski et al., 2019). For many types of tumor cells, the capability to degrade ECM by matrix metalloproteinases (MMPs) enhances their mobility (Wolf et al., 2013). However, CTLs migrate in an MMP-independent manner in collagen I due to their lack of collage I degrading-MMPs (Edsparr et al., 2011; Jablonska-Trypuc et al., 2016).

For migrating human CTLs, a positive correlation between nuclear deformation and cell speed in 3D collagen matrices was observed. The nuclei of cells are adopted hourglass-like deformation in migrating CTLs, very likely through confined spaces. The diameters of the neck of hourglass (cross-section) decreases with enhanced density of collagen. The nucleus is the most rigid intracellular organelle, which provides protection of the chromatin content (Feng and Kornmann, 2018; Lammerding, 2011). As reported in many cell types, severe deformation or even rupture of nuclei leads to DNA damage and ultimately cell death (Denais et al., 2016). In dense ECM, the enhanced nuclear deformation could therefore also lead to an elevated level of CTL apoptosis, which could eventually also contribute to dense ECM-impaired CTL killing efficiency.

Interestingly, we observed that the microtubule network is located in vicinity to the envelope of nucleus and that disruption of microtubule polymerization further enhances nuclear deformation/rupture consequently resulting in more CTL apoptosis, indicating a protective role of the microtubule network on nuclear morphology and integrity of chromatins. Compelling evidence shows that the nuclear envelope protein lamin-A/C acts as a critical structural element required for nuclear membrane organization and stability (Goldberg et al., 2008). It is also reported that nucleus morphology is associated with the microtubule network and the microtubule network could protect nucleus in coordination with Lamin A/C (Tariq et al., 2017). It is reported that Lamin A/C is induced upon activation of T cells (Rocha-Perugini and Gonzalez-Granado, 2014). Notably, microtubules are 300 fold more rigid than actin filaments (Gittes et al., 1993). Thus, microtubule network could provide a rigid protective network surrounding nucleus to prevent chromotins from being ruptured, in collaboration with Lamin A/C and/or other nuclear envelope proteins.

While CTL going through a physical restriction, our live-cell imaging shows that actin-driven protrusions leads the way through the confinement followed by translocation of the microtubule network along with the nucleus through the confined space. It is reported that in cells, deficiency in actomyosin-based contractility by myosin IIA depletion or inhibition of ROCK impedes contraction of the cell rear and fails to propel the nucleus through a restricted space (Lammermann et al., 2008; Ren et al., 2004; Wolf et al., 2013). The nucleus serves as a mechano-limiting factor, which determines the possibility of cells to go through physical confinement (Hons et al., 2018; Renkawitz et al., 2019). In our work, we found that abrogation of myosin IIA or ROCK significantly limited CTL migration in 3D collagen matrices ranging from low to intermediate-density. It is very likely due to the absence of actomyosin contraction resulting in failing translocation of nucleus through the physical restriction. Unexpectedly, disruption of microtubule polymerization with nocodazole promotes both nuclear deformation and CTL migration in low, intermediate and high density. It indicates that the microtubule network serves as an additional mechano-limiting factor in addition to nucleus to control CTL migration in 3D, especially through restricted space (Thiam et al., 2016). In comparison, in large pores/channels, the microtubule network is not the rate-limiting factors as one recent report shows that nocodazole has no effect on T cell migration through micro-channels with width equals to or beyond 6 μm (Park and Doh, 2015), which already exceeds physical constrains in low-density collagen, porosity of which ranges from 2-5μm (Wolf et al., 2013). Integrity of both actin-cytoskeleton and the microtubule network is pivotal to execute CTL killing processes. Actin-cytoskeleton has two compensatory roles. On one hand, functionality of cortical actin is essential for TCR triggered release of lytic granules to induce destruction of target cells (Lyubchenko et al., 2003). On the other hand, recovery of cortical actin in CTLs at the contact site with the target cell terminates release of lytic granules (Ritter et al., 2017). In addition, latrunculin-A treatment for target cells could also reduce target lysis-induced by perforin (Basu et al., 2016). In our work, we show that disassembly of F-actin significantly diminishes CTL motility and killing efficiency in 3D matrices. The concentration of latrunculin-A we used was 50 μM, which should only partially disassemble F-actin as CTLs could still migrate and kill under this condition. The impairment of CTL motility and the consequent reduction in searching efficiency is the primary factor for the reduced killing by disassembly of F-actin. The effect of latrunculin-A on target cells might contribute to reduced killing to some extent, if so only as a secondary factor.

In terms of the microtubule network, re-orientation of MTOC to the immunological synapse (IS) is a hallmark for CTL activation upon recognition of target cells, which plays a key role in enriching lytic granules towards the IS (Ritter et al., 2017; Yi et al., 2013). Perturbation of the microtubule architecture in CTLs results in reduced killing efficiency but does not affect degranulation (Tamzalit et al., 2020). In our work, although the microtubule network was disrupted by nocodazole (10 μM) to a large extent, the remaining was sufficient to support lytic granule release as shown in Supplemental Fig. 8a. More importantly, the enhancement in migration of nocodazole-treated CTLs in dense ECM leads to more conjugation and a consequently elevated efficiency of target destruction. Therefore, we conclude that in dense ECM, CTL motility serves as a rate-limiting factor for killing. Several microtubule inhibitors are applied as chemotherapeutic reagents, such as vinblastine and vincristine. Interestingly, we confirm that vinblastine indeed enhances CTL-mediated elimination of tumor cells in dense ECM. Our findings suggest that these microtubule-inhibiting chemotherapeutic reagents do not only have a direct effect on abrogation of tumor cell proliferation, but also have the potential to enhance CTL killing efficiency against tumor cells in densely packed tumor microenvironment.

## Author contributions

R.Z. designed and performed most experiments and all the corresponding analyses if not mentioned otherwise; X.Z. stained collagen matrices and imaged the structure; E.S.K. carried out rheology experiments and A.d.C.B. helped interpret the results; W.Y. helped with flow cytometry; K.F. and E.C.S. established MART1-specific T cells clones; E.C.S. established NALM-6-pCaspar cell lines; M.H. and D.A. provided expertise in pCasper-pMax and the corresponding analysis; B.Q. generated concepts, designed experiments, and wrote the manuscript; all authors contributed to the writing of the manuscript and provided advice.

## Acknowledgments

We thank the Institute for Clinical Hemostaseology and Transfusion Medicine for providing donor blood; Carmen Hässig, Cora Hoxha, Gertrud Schwär and Susanne Renno for excellent technical help; Sylvia Zöphel for helping with MART1-specific T cell clones; Carsten Kummerow for technical help in ImageXpress and light-sheet microscopy. This project was funded by the Deutsche Forschungsgemeinschaft (SFB 1027; Forschungsgroßgeräte (GZ: INST 256/423-1 FUGG to M.H.), Bundesministerium für Bildung und Forschung (BMBF, 031L0133 to M.H), University of Saarland HOMFORexzellent grant (to R.Z.), and by the Leibniz-Gemeinschaft (INM Fellow to B.Q.). The authors declare no competing financial interests.

## Ethical considerations

Research carried out for this study with healthy donor material (leukocyte reduction system chambers from human blood donors) is authorized by the local ethic committee (declaration from 16.4.2015 (84/15; Prof. Dr. Rettig-Stürmer)).

## Supplementary Figures

**SupFig.1.**
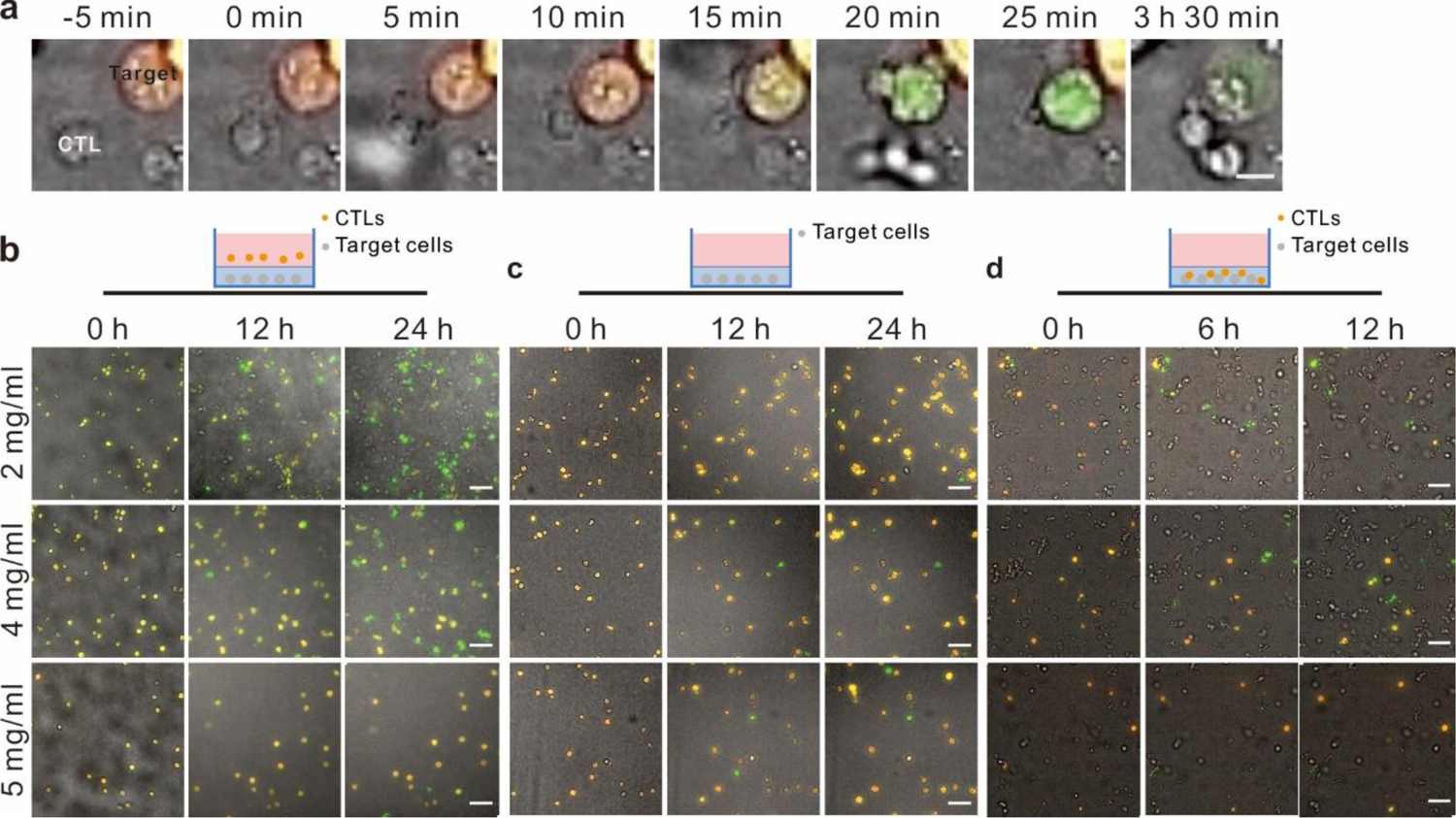
Characterization of target lysis in 3D by CTLs. **a** Determination of apoptosis versus necrosis using pCasper-expressing target cells. Target cells (SEA/SEB pulsed NAML-6-pCasper) were embedded with CTLs. Images were acquired using ImageXpress for 12 hours. The time 0 was determined when CTL contacts with the target cell. Target cells in yellow-orange were alive as indicated by FRET signal. When the target cell was undergoing apoptosis, the fluorescence was switched to green, which is pointed with white arrows. In necrotic target cells, all the fluorescence would go down abruptly, which is pointed with the yellow arrow. The scale bar is 10 µm. Results are from 3 donors. **b** Time-lapse for 3D killing. Target cells (SEA/SEB pulsed NAML-6-pCasper) were embedded in collagen in absence (**b**, **c**, Target cell only) or presence of CTLs (**d**, Target cell + CTL) in half-area 96-well plates. In **b**, CTLs were added into the wells after polymerization of collagen. Images were acquired using ImageXpress for 24 h (**b**, **c**) or 12 h (**d**). Scale bars are 50 µm. Results are from 3 donors (in **b**, **c**) or 5 donors (in **d**).

**SupFig2.**
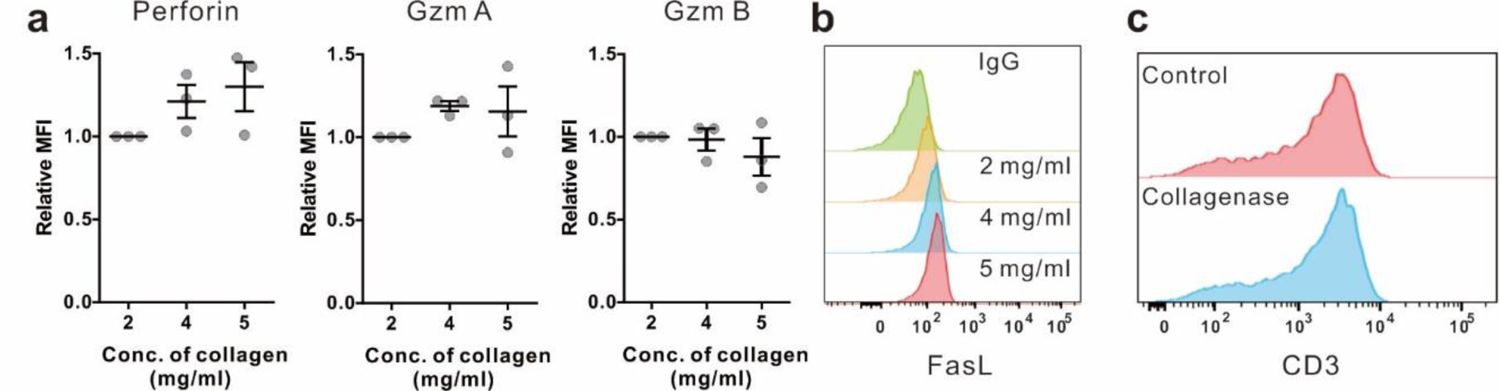
Impact of collagen density on lytic granule pathway. CTLs were embedded in collagen matrices with indicated densities and were kept at 37°C with 5% CO_2_ for 5 h (**a, b**) Then collagen matrices were digested by collagenase I for 7 min at 37°C. **a** Expression of perforin, granzyme A (GzmA), or granzyme B (GzmB) was stained with respective antibodies in fixed and permeabilized CTLs, and measured with flow cytometry (from 3 donors). **b** FasL expression was stained FasL antibody in fixed and permeabilized CTLs, and determined with flow cytometry (from 2 donors). **c** Collagenase I does not degrade surface protein on T cells. Jurkat T-cells were treated with or without collagenase for 7 min at 37°C, followed by fixation and CD3 staining without permeabilization. The samples were analyzed with flow cytometry. In **a**, one dot represents one donor. P values were assessed using the unpaired Student’s t-test. Results are presented as Mean±SD.

**SupFig3.**
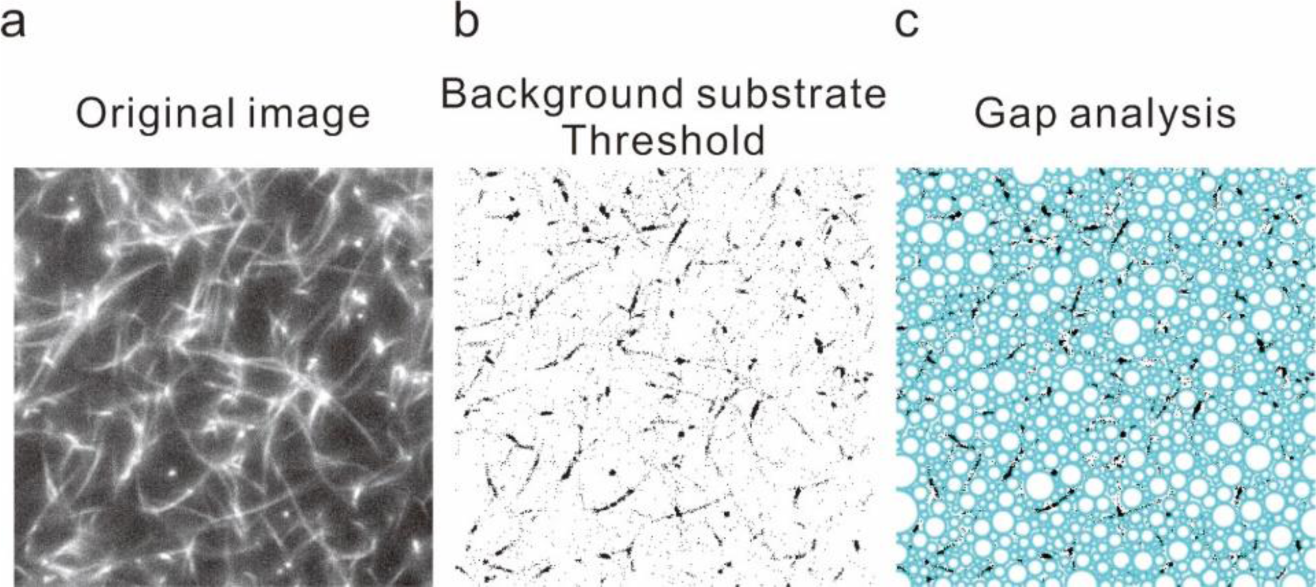
Structure of type I collagen. **a** Visualization of the collagen structure of Type I collagen (2 mg/ml) from bovine was stained with Atto 488 NHS ester, and visualized using light-sheet microscopy with a 20× objective. As elaborated in Methods, the background was subtracted (**b**), and the pores were identified by the Fiji ImageJ (BIOP version) with Max Inscribed Circles plugin (**c**), from which diameters of pores were quantified.

**SupFig4.**
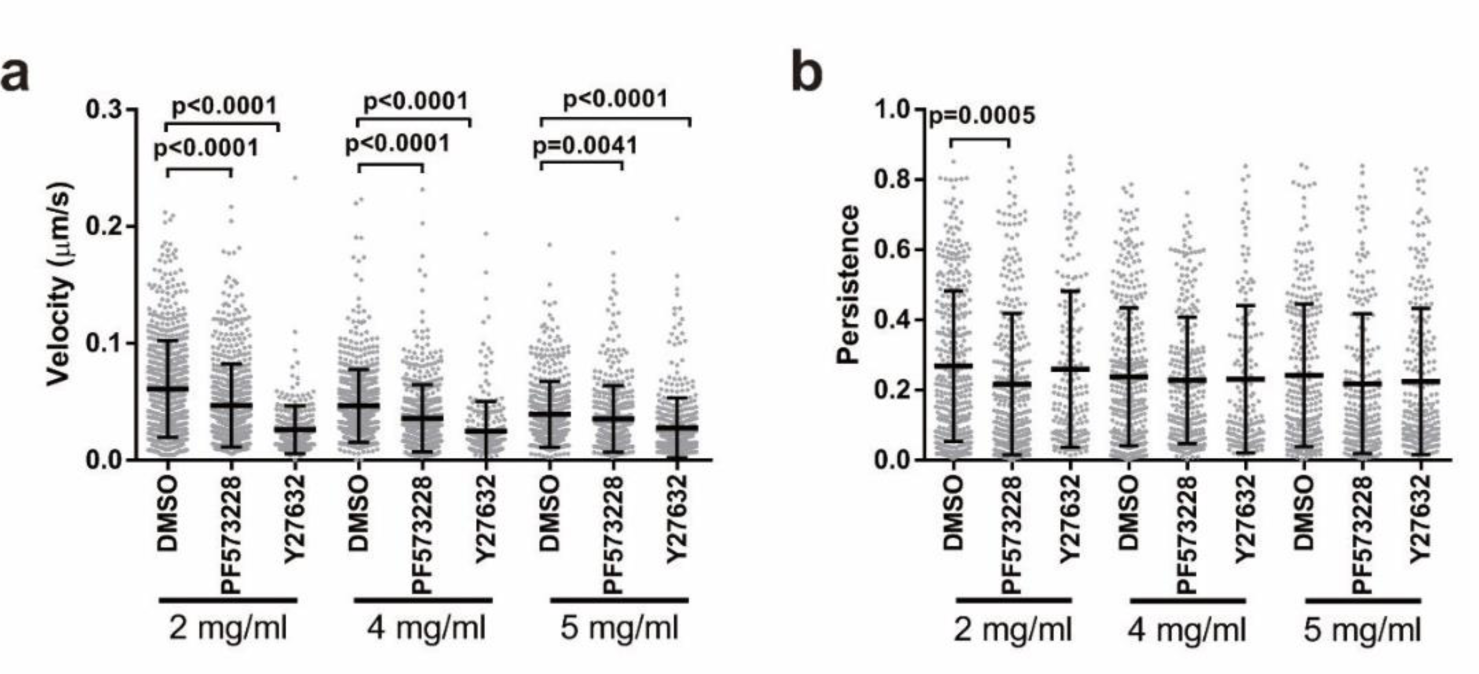
FAK and ROCK are essential for CTL migration in 3D. CTLs were embedded in planar collagen matrices with indicated densities in the presence of the correspondent inhibitor or DMSO and were visualized at 37°C with 5% CO_2_ for 30 min with an interval of 30 sec using Cell Observer (10× objective). CTLs were tracked automatically by Imaris 8.1.2. FAK inhibitor PF573228 (20 µM) and ROCK inhibitor Y27632 (10 µM) were applied. One dot represents one cell. Results are presented as Mean±SD. The results are pooled from 3 donors. P values were accessed with the Mann-Whitney test.

## Supplementary Movies

**Movie 1. CTL migration trajectories measured by lightsheet microscopy in 2 mg/ml collagen matrices.** Nuclei were labeled with overexpression of Histone 2B-GFP. Migration was visualized using light-sheet microscopy (20× objective) at 37°C for 30 min with an interval of 30 sec. Nuclei movement was tracked with spot tacking function of Imaris 8.1.2. Scale bar is 40 µm.

**Movie 2. CTL migration trajectories measured by lightsheet microscopy in 4 mg/ml collagen matrices.** Nuclei were labeled with overexpression of Histone 2B-GFP. Migration was visualized using light-sheet microscopy (20× objective) at 37°C for 30 min with an interval of 30 sec. Nuclei movement was tracked with spot tacking function of Imaris 8.1.2. Scale bar is 40 µm.

**Movie 3. CTL migration trajectories measured by lightsheet microscopy in 5 mg/ml collagen matrices.** Nuclei were labeled with overexpression of Histone 2B-GFP. Migration was visualized using light-sheet microscopy (20× objective) at 37°C for 30 min with an interval of 30 sec. Nuclei movement was tracked with spot tacking function of Imaris 8.1.2. Scale bar is 40 µm.

**Movie 4. Nuclei shape dynamics of CTLs during migration in 2 mg/ml collagen.** CTLs transfected with Histone 2B-GFP (to label nuclei, green) and LifeAct-mRuby (to label actin, red) were embedded in collagen. Migration was visualized using light-sheet microscopy (20× objective) at 37°C for 30 min with an interval of 30 sec. Nuclei dynamics were tracked with surface tacking function of Imaris 8.1.2. Scale bar is 15 µm.

**Movie 5. Nuclei shape dynamics of CTLs during migration in 4 mg/ml collagen.** CTLs transfected with Histone 2B-GFP (to label nuclei, green) and LifeAct-mRuby (to label actin, red) were embedded in collagen. Migration was visualized using light-sheet microscopy (20× objective) at 37°C for 30 min with an interval of 30 sec. Nuclei dynamics were tracked with surface tacking function of Imaris 8.1.2. Scale bar is 15 µm.

**Movie 6. Nuclei shape dynamics of CTLs during migration in 5 mg/ml collagen.** CTLs transfected with Histone 2B-GFP (to label nuclei, green) and LifeAct-mRuby (to label actin, red) were embedded in collagen. Migration was visualized using light-sheet microscopy (20× objective) at 37°C for 30 min with an interval of 30 sec. Nuclei dynamics were tracked with surface tacking function of Imaris 8.1.2. Scale bar is 10 µm.

**Movie 7. Microtubule network and F-actin dynamic during CTL migration in 4 mg/ml collagen matrices.** CTLs were transfected with EMTP-3×GFP (to label microtubules, green) and LifeAct-mRuby (to label acin, red). Migration was visualized using light-sheet microscopy (20× objective) at 37°C for 30 min with an interval of 6 sec. Scale bar is 7 µm.

